# Calling Somatic SNVs and Indels with Mutect2

**DOI:** 10.1101/861054

**Authors:** David Benjamin, Takuto Sato, Kristian Cibulskis, Gad Getz, Chip Stewart, Lee Lichtenstein

## Abstract

Mutect2 is a somatic variant caller that uses local assembly and realignment to detect SNVs and indels. Assembly implies whole haplotypes and read pairs, rather than single bases, as the atomic units of biological variation and sequencing evidence, improving variant calling. Beyond local assembly and alignment, Mutect2 is based on several probabilistic models for genotyping and filtering that work well with and without a matched normal sample and for all sequencing depths.

## Introduction

To understand the genetics of cancer, we must accurately detect somatic mutations. Due to such factors as contaminating normal cells, subclonality, and copy number variations, somatic mutations may have low allele fractions. Such mutations are difficult to distinguish from artifacts due to sample preparation, sequencing error, and mapping error. To find somatic variants Mutect2 employs local assembly and alignment, a Bayesian somatic genotyping model, and a novel filtering scheme. As a GATK 4 tool it runs on both local and cloud-based data and has a full pipeline written in the Broad Institute’s Workflow Development Language (WDL).

## Methods

Roughly speaking, Mutect2 combines the GATK’s local assembly and pair-HMM read-to-haplotype alignment, which it shares with HaplotypeCaller [1], with somatic-specific genotyping and filtering. The discussion below will focus on methods unique to Mutect2; methods shared with HaplotypeCaller are deferred to the supplemental material.

### Finding Active Regions

Mutect2 triages sites for possible somatic variation and determines intervals over which to assemble reads by assigning each site’s read pileup a log odds for somatic activity via a simplified version of the somatic genotyping model, below. A site is considered “active” if its log odds exceed some threshold. Like HaplotypeCaller, Mutect2 chooses for assembly intervals that surround each active site with some margin. The details of this are discussed in the supplemental material.

### Assembling Haplotypes

Mutect2 shares almost all of its local assembly code with HaplotypeCaller^[1]^ Like HaplotypeCaller, Mutect2 creates a de Bruijn-like graph out of kmerized reads, prunes the graph of spurious paths, attempts to bring “dangling” paths – that is, paths that start at non-reference source vertices or end at non-reference sink vertices – back to the reference path, and sets candidate haplotypes to be the highest-scoring paths from the reference source to the reference sink, where the score is the product of branching ratios of edges in the path that come from a vertex with out-degree greater than 1. An important difference is the novel adaptive pruning algorithm we devised for Mutect2. Whereas HaplotypeCaller prunes chains – that is, maximal non-branching subgraphs – based on a constant threshold for the maximal edge multiplicity in the chain, Mutect2 uses a version of its active region likelihoods model to score chains. In this model, the role of possible variants or errors is played not by pileup elements but by branching edges. Each chain has a left likelihood ratio that considers the leftmost edge multiplicity of the chain compared to the total out-multiplicity of its first vertex, and a right likelihood ratio based on the rightmost edge multiplicity versus the total in-multiplicity of its last vertex. The model learns a global error rate in a first pass where chains are not pruned but chains with likelihood ratios below the threshold are used to determine an empirical error rate. Then in the second pass this learned error rate is used to calculate more refined likelihoods and prune chains from the graph. This adaptive pruning is especially effective for samples with high and/or non-uniform read depth, such as mitochondria, exomes, and RNA.

### Somatic Genotyping

Mutect2 uses the GATK’s implementation of the Pair-HMM probabilistic model for pairwise sequence alignment to assign a likelihood for each read to have been sequenced from each candidate haplotype. This yields a matrix of read-haplotype likelihoods for the tumor sample and, if present, the normal sample. Mutect2 converts this matrix into a fragment-haplotype matrix by adding log-likelihoods of paired reads^[2]^ These fragment-haplotype matrices are converted to fragment-allele matrices at each locus where any haplotype differs from the reference by setting a fragment’s allele likelihood to be its maximum likelihood among haplotypes exhibiting that allele. Mutect2’s somatic likelihoods model for genotyping based on a fragment-allele likelihood matrix is inherently multiallelic. Pair-HMM gives the fragment-allele likelihoods matrix *ℓ_ra_ ≡ P*(read *r*|allele *a*). We assume a latent vector **f** of allele fractions with a Dirichlet prior *P*(**f**) = Dir(**f**|***α***) and assign each fragment a latent indicator **z** such that **z**_*ra*_ = 1 if read *r* was sequenced from DNA supporting allele *a* and 0 otherwise. **f** is the prior for 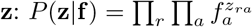. The full-model likelihood is therefore

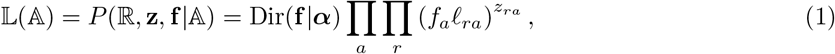

where 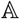 is the set of alleles. We describe the mean field variational Bayes approach to marginalizing **f** and **z** to obtain the model likelihood as a function of 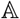 in the supplementary material.

The likelihood ratio for an allele is defined as the model evidence when the allele is excluded from 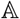, keeping all other alleles, dividing by the model evidence when 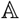 includes all alleles.

### Filtering

The companion GATK tool FilterMutectCalls filters the potential somatic variants output by Mutect2 based on the annotations that Mutect2 emits. FilterMutectCalls first estimates the probability that each candidate is a somatic variant, as opposed to a germline variant, sequencing error etc. It then chooses the probability threshold that maximizes the estimated F score - the harmonic mean of recall and precision - and filters accordingly. Error probabilities are derived from several different error models, the simplest of which are hard filters that assign an error probability of 1 when an annotation exceeds some threshold. These include filters for variants with an excess of low-quality supporting bases, poor mapping quality, support coming exclusively near the end of reads, a large mismatch in lengths between variant- and reference-supporting fragments, and local haplotypes with too many variants. FilterMutectCalls also filters sites in a “panel of normal” blacklist, where the panel of normals is obtained by running Mutect2 on a set of normal samples and blacklisting all sites that appear in more than one sample’s calls with the GATK tool CreateSomaticPanelOfNormals.

In addition to these hard filters FilterMutectCalls applies several probabilistic models for error modes that can be explicitly modeled. The germline model calculates the probability that a variant is a germline event by subjecting reads to a diploid genotyping model’^[3]^ The normal artifact model applies the somatic genotyping model to the matched normal and learns a parameter representing the extent to which this serves as a proxy for artifacts in the tumor. The contamination model uses the estimate of cross-sample contamination from the GATK tool CalculateContamination to calculate the probability that variants are due to contamination. The short tandem repeat (STR) model handles insertions and deletions that may be due to polymerase slippage. Finally, there are models for strand bias and orientation bias artifacts, the latter of which is crucial for good performance on FFPE samples.

Many of these models compute a posterior probability of error by comparing the likelihoods of some artifact against the likelihood of somatic variation. The latter depends on a somatic clustering model that determines the spectrum of tumor allele fractions and the overall somatic mutation rate. This and all other aspects of filtering are discussed in detail in the supplement.

## Results

We have validated Mutect2 against several truth data sets: the ICGC-TCGA-DREAM Somatic Mutation Calling Challenge [2] synthetic bams, in vitro mixtures of germline samples, normal sample replicates, and a large set of TCGA WES bams with matched WGS bams. All of our validations are run with default settings of Mutect2 and FilterMutectCalls, with one exception: for the in vitro mixtures we turn off the germline filter and the panel of normals because the true “somatic” mutations are really common germline variants.

### DREAM Challenge

We evaluated Mutect2 against the first four synthetic WGS tumor-normal pairs from the DREAM Challenge. These bams are from real data, with “somatic variants” spiked in in silico using BamSurgeon [3] and hence have known truth data. We ran Mutect2 and FilterMutectCalls on each pair, excluding the same masked intervals as the original DREAM challenge, and measure sensitivity and precision with respect to the synthetic truth data. That is, our metrics are identical to those of the Challenge and of its public leaderboard^[4]^. Note that challenges 1 and 2 did not contain indels. Table 1 summarizes the results from these evaluations.

**Table 1.** ICGC-TCGA-DREAM Somatic Mutation Calling Challenge.

| Challenge | Total true variants | Sensitivity | Precision |
| --- | --- | --- | --- |
| DREAM 1 SNV | 3281 | 0.975 | 0.954 |
| DREAM 2 SNV | 3979 | 0.979 | 0.944 |
| DREAM 3 SNV | 6391 | 0.929 | 0.966 |
| DREAM 3 indel | 6494 | 0.897 | 0.976 |
| DREAM 4 SNV | 11826 | 0.843 | 0.959 |
| DREAM 4 indel | 10288 | 0.814 | 0.984 |

### Normal-Normal Calls

A popular way to measure the false positive rate of somatic variant callers is to assign one normal sample as a “tumor” and a technical replicate of the same sample as the “matched normal.” In this case, any calls are by definition false positives^[5]^. We ran Mutect2 and FilterMutectCalls on the 12 = 4 × (4 − 1) ordered pairs of four NA12878 exome replicates prepared and sequenced at the Broad Institute. We summarize these results below in Table 2. The total covered territory of these exomes is about 37 Mb, so a total of 7 false positives corresponds to a false positive rate of one call per 5 Mb.

**Table 2.** Total SNV, Indel false positives in normal-normal calls on NA12878 replicates.

| Tumor | Normal |  |  |  |  |  |
| --- | --- | --- | --- | --- | --- | --- |
|  | 1 | 2 | 3 | 4 | 5 | 6 |
| 1 |  | 0,0 | 0,1 | 0,0 | 2,0 | 3,2 |
| 2 | 0,0 |  | 2,0 | 1,1 | 2,0 | 2,0 |
| 3 | 0,4 | 0,0 |  | 0,1 | 0,1 | 0,1 |
| 4 | 0,0 | 0,0 | 0,0 |  | 0,0 | 1,0 |
| 5 | 0,2 | 0,1 | 0,0 | 4,3 |  | 0,1 |
| 6 | 0,1 | 0,1 | 1,1 | 0,1 | 0,1 |  |

### Normal mixtures

We prepared WES libraries from in vitro mixtures of 5, 10, and 20 samples from the 1000 Genomes Project, replicated this three times for each mixture, and sequenced the mixtures to obtain simulated subclonal tumor samples with a range of variant allele fractions. For example, a mixture of 20 samples in which one was homozygous for a variant and one was heterozygous would exhibit an allele fraction of (1 + 2)/(2 × 20) if mixing proportions were roughly equal. By calculating the mixing fractions from the average allele fraction of singleton heterozygous alleles and obtaining germline genotypes from HaplotypeCaller, we were able to generate a large set of variants with known allele fractions. In order to avoid artifacts in our truth data we excluded variants with a population allele frequency of less than 0.001 in gnomAD. We thereby obtained a truth set of confident variants from which we could measure the sensitivity of Mutect2, although by excluding some variants from the truth set we lose the ability to measure precision. We then ran Mutect2 on all replicates of each mixture and aggregated the results by bins of predicted allele fraction and depth. We plot these aggregated sensitivity results for SNVs and indels in Figures 1

**Figure 1.**
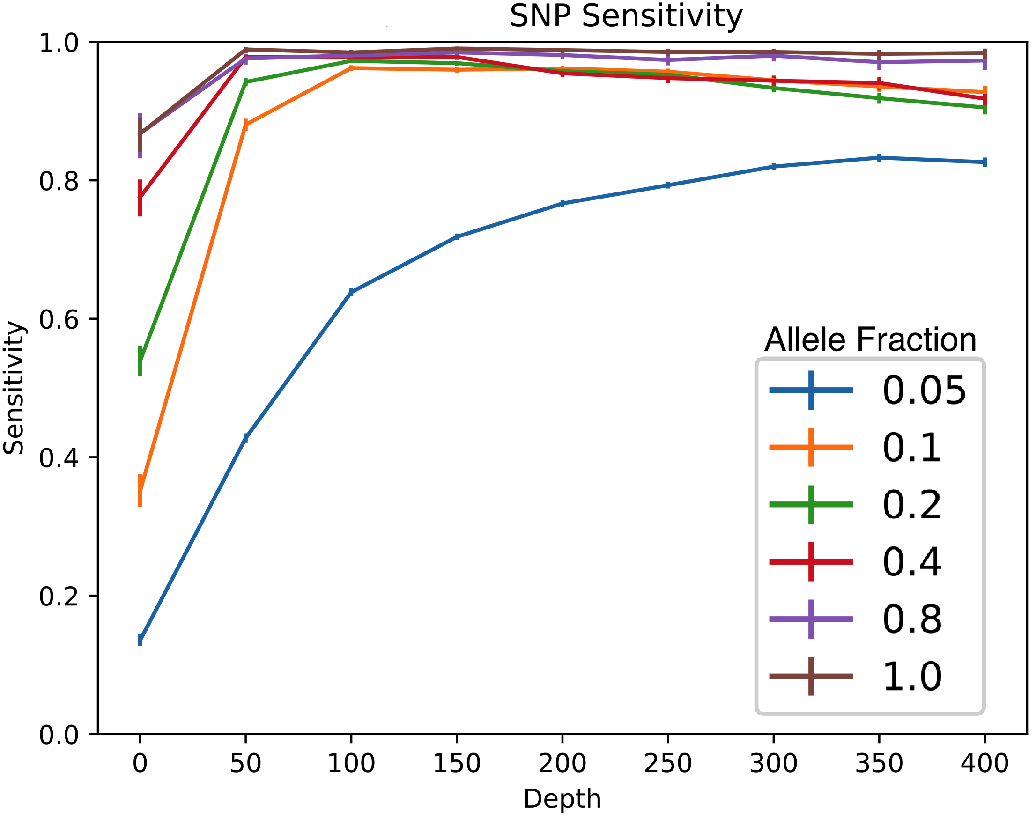
SNV sensitivity in mixtures of normal samples.

### TCGA WGS

Our final validation is based on the MC3 [4] dataset of TCGA calls. We used tumor-normal pairs from the MC3 analysis that had both WES and WGS sequencing, excluding a handful of pairs that had undergone whole-genome amplification or were marked as the non-preferred pair (eg metastasis instead of primary tumor).

**Figure 2.**
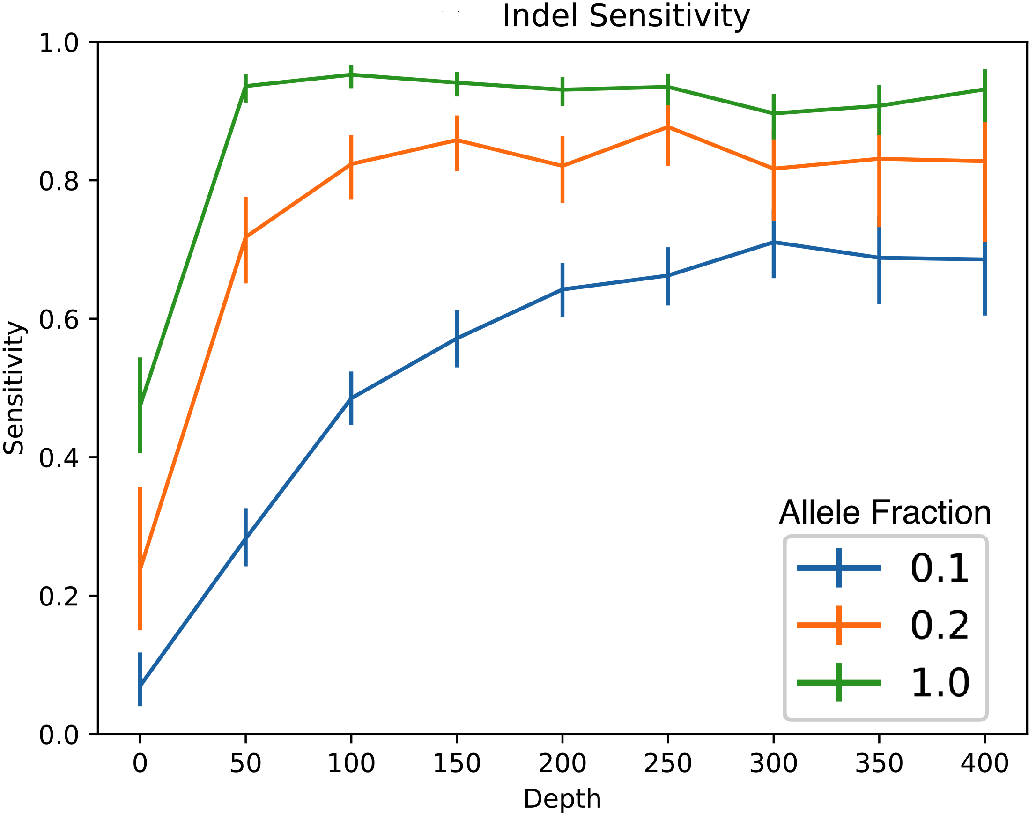
Indel sensitivity in mixtures of normal samples.

For each pair, we converted the MC3 MAF into a VCF annotated by which caller had called each variant. We ran Mutect2 and merged all Mutect2 calls, filtered and unfiltered, into the MC3 VCF to create a maximal set of candidate variations. It is expected that this union of all callers’ outputs, which includes filtered Mutect2 calls with low tumor log odds, contains essentially all true mutations. To determine which of these calls are true, we validated against the matched WGS using the GATK 4 tool ValidateBasicSomaticShortMutations, requiring at least two reads to confirm a variant. We also calculated the power for the WGS to confirm a variant, which is important for low-allele fraction mutations. If a variant had sufficient power (greater than 0.8) and was not seen in the WGS pair we counted it as a false positive. If it was not seen in the WGS pair and was underpowered, we counted it as undetermined. If it was seen in the WGS pair, regardless of power we counted it as a true positive. Then for each caller and for each pair we calculated the sensitivity and precision against this validated truth data set. The callers against which we compared Mutect2 were those in the MC3 dataset: VarScan 2 [5], SomaticSniper [6], Mutect [7], RADIA [8], and MuSE [9] for SNVs and Pindel [10], VarScan 2 [5], and Indelocator [11] for indels.

So far we have performed this analysis on 65 samples with both WES and WGS tumor-normal pairs. In subsequent versions of this manuscript we will extend the analysis to 929 samples^[6]^ We have also observed that although the WGS samples have relatively long read lengths of 150 bases they still exhibit a significant number of mapping errors in the truth data. Since Mutect2 is more aggressive about mapping error than all other tools this tends to systematically penalize the sensitivity of Mutect2 and overestimate the precision of other callers. We are working on a rigorous justification of these informal observations.

**Figure 3.**
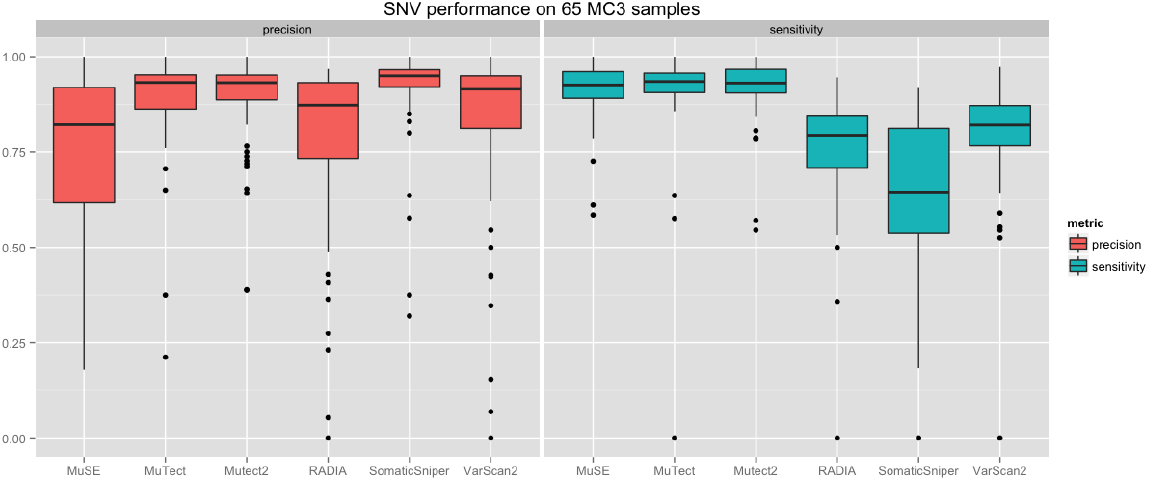
SNV sensitivity and precision of six callers on MC3 dataset.

**Figure 4.**
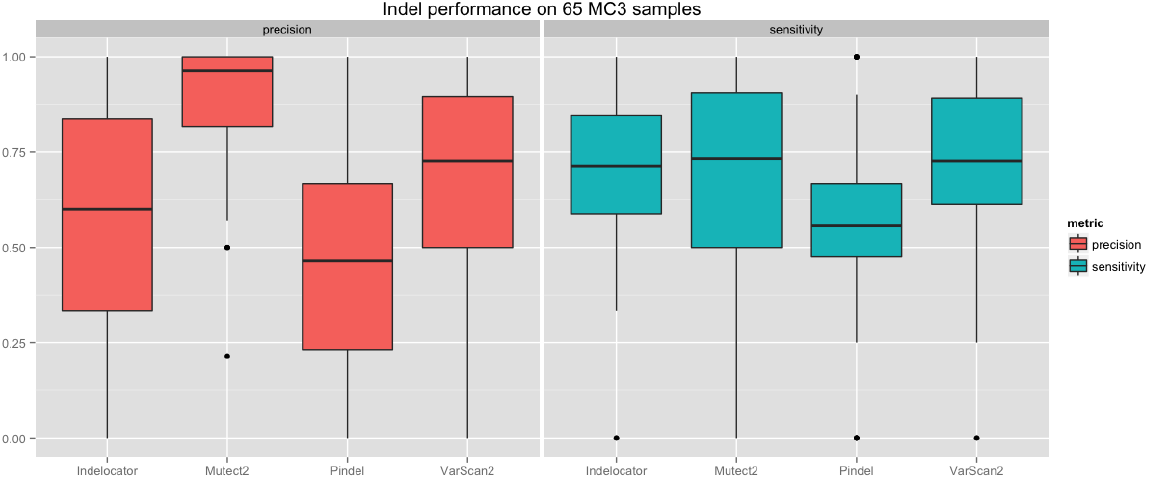
Indel sensitivity and precision of four callers on MC3 dataset.

## Acknowledgements

We acknowledge Megan Shand for discussion of mitochondria mode and adaptive pruning, Sarah Calvo for sharing unpublished data and identifying missed calls in mitochondria, Gordon Saksena for help with the MC3 dataset, and Heng Li for advice about BWA-MEM for use in the realignment filter.

## Additional Files

Additional file 1 — Supplement

Additional information on methods as well as command line invocation of Mutect2 and FilterMutectCalls.

## Footnotes

[1] Since this this has not been published elsewhere, we document the assembly of both Mutect2 and HaplotypeCaller in great detail in the supplementary material.

[2] This treats errors in paired reads as independent, which is correct for sequencing error. To account for possible shared PCR error Mutect2 caps the quality scores of bases where paired reads overlap so that their sum is bounded by an effective PCR quality score.

[3] By using the optional tumor segmentation input generated by the GATK tool CalculateContamination, this diploid model can account for copy-number variation. Otherwise it assumes that het allele fractions equal 1/2.

[4] Mutect2 performs at or near the top of each leaderboard, although note that teams were allowed an unlimited number of submission after which they saw their performance each time. Thus a tool run with default settings is at a very significant handicap.

[5] Somatic mosaicism in normal samples and mutation in culture account for some true positives but these generally occur at undetectably low allele fractions and at a negligible rate compared to callers’ error rates.

[6] Due to an unforeseen unavailability of TCGA data on the cloud for several months we have not yet been able to do this.

