## Supplemental Material for "Calling Somatic SNVs and Indels with Mutect2"

### Notes on Mutect2

David Benjamin,* Takuto Sato, and Lee Lichtenstein
*Broad Institute, 415 Main Street, Cambridge, MA 02142*
(Dated: November 30, 2019)

#### I. RUNNING MUTECT2

The following assumes that one has set up GATK 4 according to the instructions at https://github.com/broadinstitute/gatk.

##### A. Mutect2

Code block 5 shows how to invoke `Mutect2` using the gatk launch script, with optional arguments inside square braces. Note that `Mutect2` supports joint calling on an arbitrary number of tumors and matched normals, but they must be from a single individual.

```
gatk Mutect2 -R reference.fasta \
   [-L intervals.interval_list] \
   -I tumor1.bam \
   # Mutect2 may input more tumor samples from the same individual
   [-I tumor2.bam -I tumor3.bam . . .]  \
   # Mutect2 may input matched normals from the same individual
   [-I normal1.bam -I normal2.bam . . .]  \
   # For most purposes Mutect2 should be supplied with gnomAD
   [-germline-resource af-only-gnomad.vcf] \
    # Mutect2 may input a panel of normals to help identify technical artifacts
   [-pon panel_of_normals.vcf ]  \
   # Mutect2 may output orientation bias data
   # for LearnReadOrientationModel
   [--f1r2-tar-gz f1r2.tar.gz] \
   -O unfiltered.vcf
```

Listing 1: Mutect2 command

The optional germline resource can be any VCF that contains an `AF` (population allele frequency) INFO field. The Broad Institute provides a version of gnomAD stripped of all fields except `AF`. The Broad Institute also provides several panels of normals, but users with a large number (at least 50 or so) may benefit from generating their own panel with `CreateSomaticPanelOfNornals`, described below. The `--f1r2-tar-gz` argument specifies an optional output file used to filter orientation bias artifacts, such as those pervasive in FFPE samples.

##### B. FilterMutectCalls

The unfiltered output of `Mutect2` should always be passed to `FilterMutectCalls`, which emits a vcf containing the same variants, annotated with the filters that they fail, if any.

```
gatk FilterMutectCalls -V unfiltered.vcf \
   # FilterMutectCalls may input segmentation for one or more
   # tumor samples from CalculateContamination
   [--tumor-segmentation segments1.table] \
   [--tumor-segmentation segments2.table] \
```

*Electronic address:

```
 # FilterMutectCalls may input contamination estimates for one or more
 #tumor samples from CalculateContamination
[--contamination-table contamination1.table] \
[--contamination-table contamination2.table] \
# orientation bias priors from LearnReadOrientationModel
[--ob-priors priors.tar.gz] \
-O filtered.vcf
```

Listing 2: FilterMutectCalls command

The optional inputs from the GATK 4 tools `CalculateContamination` and `LearnReadOrientationModel` are described below. The `-tumor-segmentation` and `-contamination-table` arguments should be specified once per tumor sample, while there is a single (optional) `-ob-priors` argument regardless.

##### C. Creating a Panel of Normals

Currently, a panel of normals is simply a vcf of blacklisted sites flagged as recurrent artifacts[1]. One generates it by running `Mutect2` in tumor-only mode on a large number of normal samples, then running the following workflow:

```
gatk GenomicsDBImport -R reference.fasta -L intervals.interval_list \
   --genomicsdb-workspace-path pon_db \
   -V normal1.vcf \
   -V normal2.vcf \
   -V normal3.vcf . . .

gatk CreateSomaticPanelOfNormals -R reference.fasta -V gendb://pon_db -O pon.vcf
```

Listing 3: CreateSomaticPanelOfNormals command

##### D. Calculating Contamination

`CalculateContamination` is the GATK's fast, simple, and accurate method for calculating the contamination of a sample. This methods does not require a matched normal, makes no assumptions about the number of contaminating samples, and remains accurate even when the sample has a lot of copy number variation. The inputs are a bam file and a vcf of common variants – for example ExAC, gnomAD, or 1000 Genomes – with their allele frequencies. To calculate the cross-sample calculation of a tumor sample, one runs the GATK tools `GetPileupSummaries` and `CalculateContamination` as follows:

```
gatk GetPileupSummaries -I tumor.bam \
   -V common-biallelic.vcf \
   -L common-biallelic.vcf \
   -O tumor.pileups

# if a normal is present, it is helpful to obtain its pileup sumaries
gatk GetPileupSummaries -I normal.bam \
   -V common-biallelic.vcf \
   -L common-biallelic.vcf \
   -O tumor.pileups

gatk CalculateContamination -I tumor.pileups \
   # the normal pileups are useful but optional
   [-matched normal.pileups] \
   -O contamination.table \
   # it is highly recommended to produce segments for FilterMutectCalls
```

[1] `CreateSomaticPanelOfNormals` emits more information than this, but `Mutect2` does not yet use it.

```
    [-tumor-segmentation segments.table]
```

Listing 4: CalculateContamination command

Note that both `CalculateContamination`'s primary output table and its optional `-tumor-segmentation` output are recommended inputs to `FilterMutectCalls`.

##### E. Orientation Bias

The read orientation model is implemented in `LearnReadOrientationModel`, which produces the `--ob-priors` input to `FilterMutectCalls`. As mentioned above, the raw input for codeLearnReadOrientationModel is generated by specifying `--f1r2-tar-gz f1r2.tar.gz` as an output of `Mutect2`. This is then processed as follows:

```
gatk LearnReadOrientationModel \
    -I f1r2.tar.gz
    -O tumor-artifact-prior.tar.gz
```

Listing 5: LearnReadOrientationModel command

The output `tumor-artifact-prior.tar.gz` is then passed as the parameter to `FilterMutectCalls` with the `--ob-artifact-priors` argument. Note that all tar.gz files shown above contain information for every tumor sample input to `Mutect2`.

##### F. Filtering Alignment Artifacts

`FilterMutectCalls` does a fairly good job detecting all sorts of errors, including mapping artifacts. When more precision is needed and a slight loss of sensitivity is acceptable, one can run the output of `FilterMutectCalls` through `FilterAlignmentArtifacts`. This tool looks at every alt-supporting read together with its mate and maps both to the best reference available[2] using an embedded BWA-mem with parameters optimized for detecting multimapping. The tool requires a BWA mem reference image index file, which is available from the GATK resource bucket. The first step below, in which this image is generated, is usually not necessary.

```
gatk BwaMemIndexImageCreator -I reference.fasta -O reference.fasta.img

gatk FilterAlignmentArtifacts \
    # the reference argument is the bam's original alignment
    -R original-reference.fasta \
    # filtered calls from FilterMutectCalls
    -V filtered.vcf \
    -I tumor.bam \
    --bwa-mem-index-image reference.fasta.img \
    -O realignment-filtered.vcf
```

Listing 6: FilterAlignmentArtifacts command

##### G. Mitochondria mode

For mitochondrial calling one should add the `--mitochondria-mode` flag to the `Mutect2` command line. This switches several defaults from values appropriate to somatic variant calling to values that reflect the greater density of mitochondrial mutations. It also activates annotations relevant to alignment artifacts involving nuclear paralogs to mitochondrial DNA.

---

[2] That is, if the reads come from an hg19-aligned bam, they will still be realigned to hg38 if that is provided.

##### H. Force calling

One can force `Mutect2` to assemble and genotype all the variants in force-calls.vcf by adding `-alleles force-calls.vcf` to the command line. This injects all alleles in force-calls.vcf into the assembly graph, deactivates pruning of subgraphs contained them, and forces `Mutect2` to emit them in the output vcf regardless of their evidence. In order to include even filtered alleles in force-calls.vcf, use the `--genotype-filtered-alleles` flag. Force-called alleles are emitted *in addition* to any alleles that `Mutect2` would otherwise discover. This so-called GGA mode is useful for studying known driver mutations, for monitoring tumors after chemotherapy, and for validating calls from `Mutect2` or other tools against some orthogonal sequencing data.

##### I. Scattering

`Mutect2` is often parallelized by running over different intervals on different cores. Because `FilterMutectCalls` learns models from all available tumor data is is critical to merge the unfiltered output of `Mutect2` before filtering. This can be done with the GATK tool `MergeVcfs`:

```
gatk MergeVcfs -I unfiltered1.vcf -I unfiltered2.vcf . . . -O merged.vcf
```

Listing 7: MergeVcfs command

In addition, when `Mutect2` outputs a file unfiltered.vcf it also automatically generates a table unfiltered.vcf.stats that `FilterMutectCalls` uses. For scattered jobs, one must first merge the stats files with `MergeMutectStats`

```
gatk MergeMutectStats -stats unfiltered1.vcf.stats -stats unfiltered2.vcf.stats \
    . . . -O merged.stats
```

Listing 8: MergeVcfs command

Then, the merged stats must be input explicitly to `FilterMutectCalls` via the `-stats merged.stats` argument.

#### II. MUTECT2 METHODS

##### A. Finding Active Regions

`Mutect2` triages sites based on their pileup at a single base locus. If there is sufficient evidence of variation `Mutect2` proceeds with local reassembly and realignment. As in the downstream parts of `Mutect2` we seek a likelihood ratio between the existence and non-existence of an alt allele. Instead of obtaining read likelihoods via Pair-HMM, we assign each base a likelihood. For substitutions we can simply use the base quality. For indels we assign a heuristic effective quality that increases with length.

We use a version of the somatic likelihoods model, described below, with the following modifications:

- We treat every ref base as completely certain. That is, $\ell_{r,\mathrm{ref}} = 1$ for ref bases.
- We combine all alt alleles and reads supporting them into a single effective alt allele.
- Starting from the same initialization as below, we stop after a single iteration.

Under the first assumption the contribution of ref bases to the sum in Equation 9 vanishes – errorless ref bases contribute no entropy – so we only need to sum over alt bases. By the second assumption the likelihood of an alt read is related to its base quality (and associated error rate $\epsilon$) as $\ell_{r,\mathrm{alt}} = 1 - \epsilon_r$.

We compute Equation 9 for two possibilities; first, that an alt allele exists, second that only the ref allele exists. In the former case, Initializing $\bar{z}_{r,\mathrm{ref(alt)}} = 1$ for ref (alt) bases a single iteration for $q(\mathbf{f})$ gives $\boldsymbol{\beta} = (N_{\mathrm{alt}} + 1, N_{\mathrm{ref}} + 1)$. For alt reads $r$ we then obtain from Equation 6

$$\bar{z}_{r,\mathrm{alt}} = \frac{\tilde{f}_{\mathrm{alt}}(1-\epsilon_r)}{\tilde{f}_{\mathrm{alt}}(1-\epsilon_r) + \tilde{f}_{\mathrm{ref}}\epsilon_r}, \tag{1}$$

where $\tilde{f}_{\mathrm{ref(alt)}} = \exp\left(\psi(N_{\mathrm{ref(alt)}} + 1) - \psi(N+2)\right)$. We also have $g(\boldsymbol{\alpha}) = 0$ for a flat prior and $g(\boldsymbol{\beta}) = \ln(N+1) + \ln\binom{N}{N_{\mathrm{alt}}}$. For the ref-only case $g(\boldsymbol{\alpha}) = g(\boldsymbol{\beta}) = 0$ and Equation 9 reduces to $\sum_{\mathrm{alt\ reads}} \ln \epsilon_r$. Subtracting the variational log likelihoods for the two cases gives

$$\text{log odds} \approx -\ln(N+1) - \ln \binom{N}{N_{\text{alt}}} + \sum_{\text{alt reads}} H(\bar{z}_{r,\text{alt}}) + \bar{z}_{r,\text{alt}} \left( \ln(1-\epsilon_r) - \ln \epsilon_r \right), \tag{2}$$

where $H(x) = -x \ln x - (1-x) \ln(1-x)$ is the Bernoulli entropy.

We assign the pileup at a locus as either active or inactive based on a threshold for these log odds. The tumor pileup's likelihood ratio must exceed a threshold and, if present, the normal pileup's likelihoods must not exceed a different threshold. Then, using the machinery of the GATK shared with HaplotypeCaller, we convolve a Gaussian kernel over the profile of active and inactive loci (with scores of 1 and 0) and choose as assembly regions maximal spans of consecutive bases whose convolved activity exceeds a threshold, with some additional padding.

##### B. Local Assembly, Pair-HMM, and Realignment

These topics, which are common to `Mutect2` and `HaplotypeCaller`, are discussed in docs/local_assembly.pdf, docs/pair_hmm.pdf, and docs/variants_from_haplotypes.pdf in the gatk git repository. As a black box, whenever the evidence in the previous section suffices to trigger local assembly and realignment, we end up at each candidate variant site with one read-vs-allele log likelihood matrix $\ell$ for each sample, where $\ell_{ra}$ is the log probability of sequencing read $r$ given its base qualities and given that read $r$ is derived from a molecule exhibiting allele $a$. [3]

##### C. Handling Paired Reads

The fundamental unit of evidence in paired-end sequencing is not the read but the fragment of DNA from which two reads were sequenced. If we apply the somatic likelihoods model below directly to the read-vs-haplotype likelihoods matrix from Pair-HMM our model would in effect allow a read and its mate to come from two different biological haplotypes. To prevent this absurdity, we transform the read-vs-haplotype likelihoods matrix into a fragment-vs-haplotype likelihood matrix. The simplest approach, which `Mutect2` uses, is to multiply the likelihoods of all paired reads in a fragment (in log space, add) to obtain the fragment's likelihood. That is, $P(\text{fragment}|\text{haplotype}) = P(\text{read 1}|\text{haplotype})P(\text{read 2}|\text{haplotype})$. Multiplying likelihoods is justified as far as sequencing error is concerned, since sequencing errors on paired reads are statistically independent. There are, of course, shared covariates such as sequence context that influence errors on both reads. These, however, are the domain of `FilterMutectCalls`'s downstream filtering. The somatic likelihoods model of `Mutect2` is *only* concerned with distinguishing sequencing errors (in the narrow sense of an error that occurred on the sequencer itself) from possible somatic variants. `FilterMutectCalls` is responsible for distinguishing somatic variants from all other errors, such as those that occur during sample preparation and alignment.

Simply multiplying likelihoods in this way is not justified when paired reads overlap because while sequencing errors are independent, PCR errors are not. That is, a substitution occurring on both reads may be due to independent sequencing errors or to the amplification of a single PCR error. In order to force the possibility of PCR error into a model of independent reads, we enforce that the effective, multiplied, likelihood must not exceed a bound given by the probability of PCR error. To achieve this, at bases where read pairs overlap, `Mutect2` caps the total (that is, the sum, because qualities are measured in a logarithmic phred scale) base and indel qualities to user-defined PCR SNV and indel qualities. This can be adjusted for PCR-free protocols. For example, if the PCR quality is 40 and two reads overlap with base qualities of 25 and 30, each base quality is replaced with $40/2 = 20$. If the base qualities were both, say, 15, no adjustment would be needed because the probability of sequencing error dominates the probability of PCR error. This capping of overlapping base qualities occurs before Pair-HMM, while merging reads into fragments occurs after Pair-HMM.

---

[3] Technically, pair-HMM produces a read-vs-*haplotype* likelihood matrix, which is then "marginalized" to produce a set of read-vs-allele likelihood matrices. In the future, `Mutect2` may operate directly on this read-vs-haplotype matrix in order to exploit the biological fact that there are only a few haplotypes in any region.

##### D. Somatic Likelihoods Model

We have a set of potential somatic alleles and fragment-allele likelihoods $\ell_{ra} \equiv P(\text{fragment } r|\text{allele } a)$. We don't know which alleles are real somatic alleles and so we must compute, for each subset $\mathbb{A}$ of alleles, the likelihood that the fragments come from $\mathbb{A}$. A simple model for this likelihood is as follows: each fragment $r$ is associated with a latent indicator vector $\mathbf{z}_r$ with one-hot encoding $z_{ra} = 1$ iff fragment $r$ came from allele $a \in \mathbb{A}$. The conditional probabilities of fragments given alleles is $\ell_{ra}$. There is a latent vector $\mathbf{f}$ of allele fractions such that $f_a$ is the allele fraction of allele $a$, that is, the prior probability that any given fragment comes from allele $a$. Giving $\mathbf{f}$ a Dirichlet prior, we have a full-model likelihood

$$P(\mathbb{R}, \mathbf{z}, \mathbf{f}|\mathbb{A}) = P(\mathbf{f})P(\mathbf{z}|\mathbf{f})P(\mathbb{R}|\mathbf{z}, \mathbb{A}) = \text{Dir}(\mathbf{f}|\boldsymbol{\alpha}) \prod_a \prod_r (f_a \ell_{ra})^{z_{ra}} . \tag{3}$$

We want to marginalize the latent variables to obtain the evidence $P(\mathbb{R}|\mathbb{A})$, which we make tractable via a mean-field approximation $P(\mathbb{R}, \mathbf{z}, \mathbf{f}|\mathbb{A}) \approx q(\mathbf{z})q(\mathbf{f})$, which is exact in two limits. First, if there are many fragments, each allele is associated with many fragments and therefore the Law of Large Numbers causes $\mathbf{f}$ and $\mathbf{z}$ to become uncorrelated. Second, if the allele assignments of fragments are obvious $\mathbf{z}_r$ is effectively determinate, hence uncorrelated with $\mathbf{f}$. In the variational Bayesian mean-field formalism we have

$$q(\mathbf{f}) \propto \exp E_{q(\mathbf{z})}\left[\ln P(\mathbb{R}, \mathbf{z}, \mathbf{f}|\mathbb{A})\right] \propto \text{Dir}(\mathbf{f}|\boldsymbol{\alpha} + \sum_r \bar{\mathbf{z}}_r) \equiv \text{Dir}(\mathbf{f}|\boldsymbol{\beta}), \quad \boldsymbol{\beta} = \boldsymbol{\alpha} + \sum_r \bar{\mathbf{z}}_r \tag{4}$$

$$q(\mathbf{z}_r) \propto \exp E_{q(\mathbf{f})}\left[\ln P(\mathbb{R}, \mathbf{z}, \mathbf{f}|\mathbb{A})\right] \propto \prod_a \left(\tilde{f}_a \ell_{ra}\right)^{z_{ra}}, \tag{5}$$

where, with $\psi$ denoting the digamma function, the moments

$$\bar{z}_{ra} = E_{q(\mathbf{z})}\left[z_{ra}\right] = \frac{\tilde{f}_a \ell_{ra}}{\sum_{a'} \tilde{f}_{a'} \ell_{ra'}} \tag{6}$$

$$\ln \tilde{f}_a = E_{q(\mathbf{f})}[\ln f_a] = \psi(\beta_a) - \psi(\sum_{a'} \beta_{a'}) \tag{7}$$

are easily obtained from the categorical distribution $q(\mathbf{z})$ and the Dirichlet distribution $q(\mathbf{f})$[4]. We initialize $\bar{z}_{ra} = 1$ if $a$ is the most likely allele for fragment $r$, 0 otherwise and iterate Equations 4 and 5 until convergence. Having obtained the mean fields of $q(\mathbf{z})$ and $q(\mathbf{f})$, we use the variational approximation (Bishop's Eq 10.3) to the model evidence:

$$\ln P(\mathbb{R}|\mathbb{A}) \approx E_q\left[\ln P(\mathbb{R}, \mathbf{z}, \mathbf{f}|\mathbb{A})\right] - E_q\left[\ln q(\mathbf{z})\right] - E_q\left[\ln q(\mathbf{f})\right]. \tag{8}$$

The terms in Eq 8 all involve the standard moments mentioned above, so after a bit of algebraic cancellation we obtain

$$\ln P(\mathbb{R}|\mathbb{A}) \approx g(\boldsymbol{\alpha}) - g(\boldsymbol{\beta}) + \sum_{ra} \bar{z}_{ra}\left(\ln \ell_{ra} - \ln \bar{z}_{ra}\right), \tag{9}$$

where we define $g$ to be the Dirichlet distribution log normalization:

$$\ln \Gamma(\sum_a \omega_a) - \sum_a \ln \Gamma(\omega_a). \tag{10}$$

We now have the model evidence for allele subset $\mathbb{A}$. The `TLOD` emitted by `Mutect2` for an alt allele is the log evidence ratio of an allele set containing all alleles versus an allele set excluding that allele. That is, it is the log odds that an allele exists. When multiple tumor samples are given, `Mutect2` computes a single `TLOD` by combining all tumor fragments.

---

[4] Note that we didn't *impose* this in any way. It simply falls out of the mean field equations.

#### III. FILTERMUTECTCALLS METHODS

##### A. Filtering Architecture

`FilterMutectCalls` contains a set of filters, each of which computes an error probability for each candidate variant. The filters are divided into three categories: technical artifacts, non-somatic, and sequencing error. Roughly, we assume that different types of errors within a category are correlated, while different categories are independent. For example, whether sequencing errors cause several bases to be misread is independent of whether the DNA being read came from a contaminating sample and of whether an error during library preparation caused a base error prior to sequencing. To obtain an overall error probability `FilterMutectCalls` computes the maximum within categories and an independent product between categories. That is:

$$P(\text{error}) = 1 - (1 - \text{max artifact error prob})(1 - \text{max non-somatic prob})(1 - \text{sequencing error prob}). \tag{11}$$

`FilterMutectCalls` goes over an unfiltered vcf in three passes, two to learn any unknown parameters of the filters' models and to set a threshold on $P(\text{error})$, and one to apply the learned filters. This section describes methods for determining the threshold on error probability. One option is to simply set a fixed value $p$ on the error probability, below which a call is considered a real somatic variant. This non-default behavior can be set via `--threshold-strategy CONSTANT --initial-threshold <double>`.

By default `FilterMutectCalls` optimizes the F-score – the harmonic mean of recall and precision – as its default thresholding strategy. The `--f-score-beta <double>` command line argument can be set to change the relative weight of recall to precision. In order to optimize the F-score, `FilterMutectCalls` sorts all variants by the probability that they are errors, from least to greatest and calculates the F-score for thresholds in which the first $n$ variants pass, starting from $n = 0$ and ending at $n = N$, the total number of candidates. This is a cheap $O(N)$ computation because initially the expected number of true positive and false positive calls are both zero. When the threshold increases to admit a variant with error probability $p$, the expected number of true positive calls increases by $1 - p$ and the expected number of false positive calls increases by $p$. The total expected number of real variants is $\sum_n (1 - p_n)$. These quantities suffice to calculate recall and precision for every threshold.

`FilterMutectCalls` can also choose to maximize sensitivity subject to a maximum allowable false discovery rate using the `--threshold-strategy FALSE_DISCOVERY_RATE --false-discovery-rate <double>`. For this calculation `FilterMutectCalls` also sorts the error probabilities $p_n$ from least to greatest. If we allow the first $M$ variants to pass the expected false discovery rate is

$$\frac{1}{M} \sum_{n=1}^{M} p_n \tag{12}$$

This is a non-decreasing function of $M$ because it is the integral from 0 to $M$ of the monotonic function $p$, hence its derivative with respect to $M$ is $p_M$, which is monotonic. Thus it is easy to choose the highest $M$ such that the maximum false discovery rate is not exceeded.

##### B. Hard Filters

Several filters are hard filters that assign an error probability of 1 whenever some annotation exceeds a threshold. Here we summarize all the hard filters of `FilterMutectCalls`, the command line parameters that set their thresholds, and an explanation of the thresholded quantity.

| Filter | Threshold | Explanation |
|---|---|---|
| `clustered_events` | `max-events-in-region` | mutations sharing an assembly region |
| `duplicate_evidence` | `unique-alt-read-count` | unique insert start/end pairs of alt reads |
| `multiallelic` | `max-alt-alleles-count` | passing alt alleles at a site |
| `base_quality` | `min-median-base-quality` | median base quality of alt reads |
| `mapping_quality` | `min-median-mapping-quality` | median mapping quality of alt reads |
| `fragment_length` | `max-median-fragment-length-difference` | difference of alt and ref reads' median fragment lengths |
| `read_position` | `min-median-read-position` | median distance of alt mutations from end of read |
| `panel_of_normals` | `panel-of-normals` | presence in panel of normals |

##### C. Allele Fraction Clustering Model

Several of `FilterMutectCalls`'s probabilistic filters rely on the following graphical model for the tumor's spectrum of allele fractions of real variants. At the top of the model is a binary indicator for whether a variant is neither a technical artifact or a non-somatic mutation.

$$P(\text{consider}) = (1 - \text{max artifact error prob})(1 - \text{max non-somatic prob}). \tag{13}$$

Note that possible sequencing errors, with error probability captured by the independent reads assumption of the `Mutect2` log odds, *are* considered. Next, a candidate SNV has prior probability $\pi_{\text{real}} = \pi_{\text{SNV}}/3$ to be real and prior probability $1 - \pi_{\text{SNV}}/3$ to be a sequencing error, while a candidate indel of length $L$ (negative for deletions, positive for insertions) has prior probability $\pi_{\text{real}} = \pi_L$ to be real. Real variants are partitioned according to a mixture model with weights $\pi_H$ for a high-allele fraction cluster, $\pi_B$ for a "background cluster", and $\pi_D = 1 - \pi_H - \pi_B$ to be further clustered into according to a Dirichlet process with concentration $\alpha$. Each non-error cluster has a characteristic distribution of allele fractions: $\text{Beta}(\alpha_H, \beta_H)$ and $\text{Beta}(\alpha_B, \beta_B)$ for the high-allele fraction and background clusters, and Dirac $\delta$ functions for the Dirichlet clusters. The idea of this partitioning is that there will usually be sharply-defined allele fractions corresponding to the hets of different subclones, which are captured by the Dirichlet clusters, but that homozygosity and copy-number variation produce deviations from this clustering.

Provided that we stochastically assign the $i$th variant as non-artifactual, the prior probabilities for each possibility are described by a modified Chinese Restaurant Process scheme:

$$P(\text{Sequencing error}) = 1 - \pi_{\text{real}} \tag{14}$$

$$P(\text{High AF}) = \pi_{\text{real}} \pi_H \tag{15}$$

$$P(\text{Background}) = \pi_{\text{real}} \pi_B \tag{16}$$

$$P(\text{Dirichlet}_j) = \pi_{\text{real}} \pi_D \frac{N_j^{-i}}{N_D^{-i} + \alpha} \tag{17}$$

$$P(\text{Dirichlet}_{\text{new}}) = \pi_{\text{real}} \pi_D \frac{\alpha}{N_D^{-i} + \alpha}, \tag{18}$$

where $\text{Dirichlet}_j$ denotes the $j$th existing Dirichlet Process cluster, $\text{Dirichlet}_{\text{new}}$ denotes a newly-created Dirichlet cluster, $N_j^{-i}$ is the number of variants besides the $i$th assigned to cluster $j$, and $N_D^{-i}$ is the total number of variants besides the $i$th assigned to Dirichlet clusters.

`Mutect2` emits the likelihood ratio between sequencing error and a real variant assuming a flat prior. However, we can approximately compute the likelihood ratio that a non-flat Beta prior would have produced. We can do this because in Equation 9 a change in the prior allele fraction is unlikely to affect $\bar{z}$ very much - usually the allele that a read supports is clear regardless of prior. Therefore, only the term $g(\boldsymbol{\alpha}) - g(\boldsymbol{\beta})$ changes. Since $\boldsymbol{\beta} = \boldsymbol{\alpha} + \mathbf{n}$, where $\mathbf{n}$ is the vector of observed read counts per allele we can convert the log-likelihood obtained with flat prior $\boldsymbol{\alpha}_0$ to one reflecting an allele fraction prior $\boldsymbol{\alpha}$ by adding

$$\Delta\text{log odds}(\boldsymbol{\alpha}) = \Delta \ln P(\mathbb{R}|\mathbb{A}) = [g(\boldsymbol{\alpha}) - g(\boldsymbol{\alpha} + \mathbf{n})] - [g(\boldsymbol{\alpha}_0) - g(\boldsymbol{\alpha}_0 + \mathbf{n})]. \tag{19}$$

Since only *ratios* of likelihoods are meaningful, we may WLOG set the likelihood of reads given sequencing error to be 1 and then set the likelihood fo each allele fraction cluster to be $\exp[\text{Mutect2 log odds} + \Delta\text{log odds}(\boldsymbol{\alpha})]$. The likelihood of a new cluster is the average over all possible new allele fractions of the likelihood, which is, by definition, simply the original `Mutect2` likelihood that integrated over a flat prior. Thus we have likelihoods for every cluster. Combined with the priors, this gives us the conditional posterior for Gibbs sampling the cluster assignment of a candidate variant.

We can estimate the parameters of the model by maximum likelihood. For example, we estimate $\pi_L$ as the total number of length-$L$ indels in the non-error clusters divided by the total number of callable sites[5], $\pi_B$ as the total number of variants assigned to the background cluster divided by the total number of non-error assignments. The allele fraction of a Dirichlet cluster is estimated as its total alt read count divided by its total read count. Finally, the beta distributions of the high-allele fraction and background clusters are fit by brute force likelihood optimization.

Once the model converges, the combination of cluster weights and allele-fraction-adjusted likelihoods lets us compute the posterior probability that a candidate variant is a sequencing error, which determines the `weak_evidence`

---

[5] This is emitted by `Mutect2` and passed to `FilterMutectCalls`.

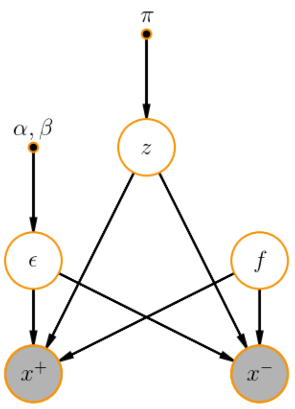


FIG. 1: The probabilistic graphical model for the strand artifact model

filter. Since the clusters also define distributions on alt allele counts – the high-AF and background clusters define beta-binomials and the Dirichlet clusters define binomial – we also have the mixture model distribution $P(\text{alt count}|\text{total count}, \text{somatic})$, which the contamination and germline filters use.

##### D. Strand Artifact Model

The strand artifact filter detects sequencing artifacts in which the evidence for the alt allele consists entirely of forward strand reads alone or reverse strand reads alone. We must detect this while taking into account the fact that at some loci, such as near the end of an exome target, *all* reads are biased towards one direction, and therefore a bias towards a particular strand among alt reads is no cause for alarm.

Let $z \in \{z_+, z_-, z_o\}$ be a latent random variable with 1-hot encoding that represents the artifact state of a suspected variant i.e. $z_+ = 1$ for a forward strand artifact, $z_- = 1$ for a reverse artifact, and $z_o = 1$ otherwise. At each locus, let $x_\pm$ be the number of forward (+) or reverse (-) strand alt reads and let $n_\pm$ be the total depth for each strand. By modeling $x_\pm$ relative to $n_\pm$ we account for inherent strand bias due to, for example, reads falling at the end of an exome target and do not confuse it for an artifact. Let $f$ be the allele fraction of true variation in case $z_o = 1$. Let $\epsilon$ be the strand bias error rate and let $\theta$ be the non-strand-biased error rate. We will ignore the case in which significant strand bias coincides with real variation, first because this is exceedingly rare and ignoring it has a negligible effect on the parameters of our model, and secondly because such variants should be considered true positives.

The conditional distributions of our model are binomial

$$x_+|\epsilon, \theta, f, z_+ = 1 \sim \text{Bin}(x_+|n_+, \epsilon) \tag{20}$$
$$x_+|\epsilon, \theta, f, z_- = 1 \sim \text{Bin}(x_+|n_+, \theta) \tag{21}$$
$$x_+|\epsilon, \theta, f, z_o = 1 \sim \text{Bin}(x_+|n_+, f), \tag{22}$$

and similarly for $x_-$. Putting beta priors on $\epsilon$, $\theta$, and $f$, with parameters $(\alpha_\epsilon, \beta_\epsilon)$, $(\alpha_\theta, \beta_\theta)$, and $(\alpha_f, \beta_f)$ and marginalizing latent parameters we obtain likelihoods

$$P(x_+, x_-|z_\pm = 1) = \text{BetaBinom}(x_\pm|n_\pm, \alpha_\epsilon, \beta_\epsilon)\text{BetaBinom}(x_\mp|n_\mp, \alpha_\theta, \beta_\theta) \tag{23}$$
$$P(x_+, x_-|z_o = 1) = \int_0^1 \text{Beta}(f|\alpha_f, \beta_f)\text{Binom}(x_+|n_+, f)\text{Binom}(x_-|n_-, f)df \tag{24}$$
$$= \frac{\binom{n_+}{x_+}\binom{n_-}{x_-}}{\binom{n_+ + n_-}{x_+ + x_-}}\text{BetaBinom}(x_+ + x_-|n_+ + n_-, \alpha_f, \beta_f) \tag{25}$$

Finally, we let $\pi/2$ be the prior probability of a forward or reverse strand artifact. From the above equations it is straightforward to calculate the posterior probability of $z$ and to learn $\pi$ iteratively via the EM algorithm. It is somewhat more complicated to learn $(\alpha_\epsilon, \beta_\epsilon)$ and $(\alpha_\theta, \beta_\theta)$, so we treat these as fixed hyperparameters. We use a flat prior $\alpha_f = \beta_f = 1$ for the true allele fraction.

##### E. Germline Filter

Suppose we have detected an allele such that its (somatic) likelihood in the tumor is $\ell_t$ and its (diploid) likelihood in the normal is $\ell_n$[6]. By convention, both of these are relative to a likelihood of 1 for the allele *not* to be found. If we have no matched normal, $\ell_n = 1$. Suppose we also have the population allele frequency $f$ of this allele. Then the prior probabilities for the normal to be heterozygous and homozygous alt for the allele are $2f(1-f)$ and $f^2$ and the prior probability for the normal genotype not to contain the allele is $(1-f)^2$. Finally, let the prior for this allele to arise as a somatic variant be $\pi$.

If the variant exists in the tumor as a real somatic variant it has likelihood $\ell_t^S$, the adjusted `Mutect2` likelihood accounting for allele fraction clustering. If it exists as a germline het in a segment with minor allele fraction $m$ the likelihood is $\ell_t(m)$ for the alt minor case and $\ell_t(1-m)$ for the alt major case, after adjusted the `Mutect2` likelihood toa ccount for pinning the allele fraction to $m$ or $1-m$. We need the unnormalized probabilities of three possibilities:

1. The variant exists in the tumor and the normal as a germline het. This has unnormalized probability $f(1-f)\ell_n(1-\pi)\left(\ell_t(m) + \ell_t(1-m)\right)$.
2. The variant exists in the tumor and the normal as a germline hom alt. This has unnormalized probability $f^2 \ell_n \ell_t(1)(1-\pi)$.
3. The variant exists in the tumor but not the normal. This has unnormalized probability $(1-f)^2 \ell_t^S \pi$.

We exclude possibilities in which the variant does not exist in the tumor sample because we really want the conditional probability that the variant is germline given that it would otherwise be called. The normalized sun of the first two possibilities is the germline error probability.

So far we have assumed that the population allele frequency $f$ is known, which is the case if it is found in our germline resource, such as gnomAD. If $f$ is not known we must make a reasonable guess as follows. Suppose the prior distribution on $f$ is $\mathrm{Beta}(\alpha, \beta)$. The mean $\alpha/(\alpha+\beta)$ of this prior is the average human heterozygosity $\theta \approx 10^{-3}$, so we have $\beta \approx \alpha/\theta$. We need one more constraint to determine $\alpha$ and $\beta$, and since we are concerned with imputing $f$ when $f$ is small we use a condition based on rare variants. Specifically, the number of variant alleles $n$ at some site in a germline resource with $N/2$ samples, hence $N$ chromosomes, is given by $f \sim \mathrm{Beta}(\alpha, \beta), n \sim \mathrm{Binom}(N, f)$. That is, $n \sim \mathrm{BetaBinom}(\alpha, \beta, N)$. The probability of a site being non-variant in every sample is then $P(n = 0) = \mathrm{BetaBinom}(0|\alpha, \beta, N)$, which we equate to the empirical proportion of non-variant sites in our resource, about 7/8 for exonic sites in gnomAD. Solving, we obtain approximately $\alpha = 0.01, \beta = 10$ for gnomAD. Now, given that some allele found by `Mutect2` is not in the resource, the posterior on $f$ is $\mathrm{Beta}(\alpha, \beta + N)$, the mean of which is, since $\beta << N$, about $\alpha/N$. By default, `Mutect2` uses this value.

##### F. Contamination Filter

Suppose our tumor sample has contamination fraction $\alpha$ and that at some site we have $a$ alt reads out of $d$ total reads. Suppose further that the alt allele has population allele frequency $f$. We estimate the probability that these alt reads came from a contaminating sample and not from a true somatic variant. Let $\pi$ be the prior probability of a somatic variation from the allele fraction clustering model. Then alt read count likelihood $P(a|d, \mathrm{somatic})$ is also given by this model. The possibility that the reads are due to contamination has prior $1-\pi$ and we take the maximum likelihood among two models of contamination.

If there are multiple contaminants we approximate each contaminant read as independent so that

$$P(a|d, \text{many contaminant}) = \mathrm{Binom}(a|d, \alpha f). \tag{26}$$

If there is a single contaminating sample it is heterozygous with probability $2f(1-f)$ and homozygous for the alt with probability $f^2$, in which cases fractions $\alpha/2$ and $\alpha$ of all reads to be alt contaminants. The contaminant is

[6] This is the total likelihood for het and hom alt in the normal.

homozygous for the ref with probability $(1-f)^2$, which yields no alt reads. Thus

$$P(a|d, \text{one contaminant}) = 2f(1-f)\text{Binom}(a|d, \alpha/2) + f^2\text{Binom}(a|d, \alpha) + (1-f)^2\text{I}[a=0]. \tag{27}$$

Taking the maximum of these as the likelihood given contamination, we can easily compute the probability of contamination error.

#### G. Read Orientation Artifact Filter

The read orientation artifact, also known as the orientation bias artifact, arises due to a chemical change in the nucleotide during library prep that results in, for example, G base-paring with A. This kind of artifact has a clear signature (e.g. C to A SNP that occurs predominantly for the middle C in the DNA sequence CCG), and it's single-stranded in nature. Downstream, this artifact manifests as low allele fraction SNPs whose evidence for the alt allele consists almost entirely F1R2 reads or F2R1 reads. A read pair is F1R2 (forward 1st, reverse 2nd) if the sequence of bases in Read 1 maps to the forward strand of the reference (F1), and the sequence of Read 2 to the reverse strand of the reference (R2). F2R1 is defined similarly.

Without loss of generality, suppose that the reference context at locus $i$ is ACT. Let $\mathbf{z}_i$ denote the genotype at locus $i$ with the one-hot encoding $z_{ik} = 1$ iff the genotype of locus $i$ is $k$, where the possible genotypes are

$$z_i \in \{\text{F1R2}_A, \text{F1R2}_G, \text{F1R2}_T, \text{F2R1}_A, \text{F2R1}_G, \text{F2R1}_T, \text{Hom Ref}, \text{Germline Het}, \text{Somatic Het}, \text{Hom Var}\}$$

$z_i = \text{F1R2}_\text{A}$ denotes that at locus $i$ we have an artifact in which the evidence for alt allele A consists entirely of reads in the F1R2 orientation. The remaining artifact states are defined analogously. Let $\pi$ denote the prior probabilities of the $\mathbf{z}_i$ under the reference context ACT. Then we have

$$P(\mathbf{z}_i) = \prod_k \pi_k^{z_{ik}} \tag{28}$$

The number of alt reads at a locus depends on the genotype $z_i$. Let $n_i$ and $m_i$ denote the total depth and alt depth at locus $i$, respectively. The conditional distribution of $m_i$ is

$$P(m_i|z_{ik} = 1) = \text{BetaBinomial}(m_i|n_i, \alpha_k, \beta_k) \tag{29}$$

where $\alpha_k$ and $\beta_k$ are fixed hyperparameters for genotype $z_k$. When the site's genotype indicates in $m_i$ alt reads we expect a heavily skewed distribution of F1R2 reads. This is captured in the conditional distribution of F1R2 alt reads. Let $c_i$ denote the number of F1R2 reads among the $m_i$ alt reads at locus $i$. Then we have

$$P(c_i|m_i, z_{ik} = 1) = \text{BetaBinomial}(c_i|m_i, \alpha'_k, \beta'_k) \tag{30}$$

We learn the prior artifact probabilities $\pi$ based on the observed values of $n_i$, $m_i$, $c_i$ for each of $N$ loci using the EM algorithm. In the E-step, we compute the posterior probabilities of $\mathbf{z}_i$ for $i = 1...N$. The joint probabilities of $\mathbf{z}$ factorizes over $i$, thus the posteriors over $\mathbf{z}$ are independent across loci.

$$P(z_{ik} = 1|m_i, c_i) \propto P(z_{ik} = 1, m_i, c_i) = \pi_k \text{BetaBinomial}(m_i|n_i, \alpha_k, \beta_k)\text{BetaBinomial}(c_i|m_i, \alpha'_k, \beta'_k) \tag{31}$$

In the M-step we maximize the expectation of the log complete-data likelihood with respect to $\pi$. The log complete data likelihood is given as

$$\ln P(\mathbf{z}, \mathbf{m}, \mathbf{c}) = \sum_i \sum_k z_{ik} \big( \ln \pi_k + \ln \text{BetaBinomial}(m_i|n_i, \alpha_k, \beta_k) + \ln \text{BetaBinomial}(c_i|m_i, \alpha'_k, \beta'_k) \big) \tag{32}$$

Maximizing the log likelihood under the constraint $\sum_k \pi_k = 1$ gives us

$$\pi_k = \frac{N_k}{N} \tag{33}$$

where $N_k = \sum_i P(z_{ik}|m_i, c_i)$ is the effective count of loci with genotype $k$. We alternate E-step and M-step until convergence. We then use the learned prior genotype probabilities to compute the posterior artifact probabilities of variants in a vcf. The filtering threshold is set such that the false discovery rate doesn't exceed a specified value, as described below.

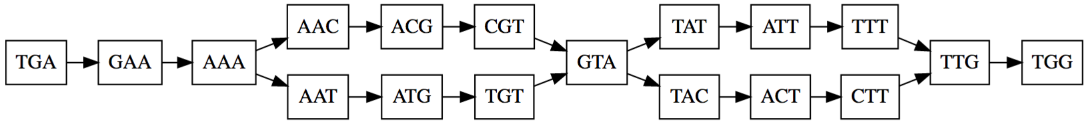


FIG. 2: Two haplotypes yielding assembly graph with two bubbles and four paths due to too-small $k$.

##### H. Polymerase Slippage

For indels in short tandem repeats (STRs) `FilterMutectCalls` uses a simple model for the possibility that alt reads are due to polymerase slippage. The prior $\pi_L$ for a real variant of length $L$ comes from the allele fraction clustering model. `FilterMutectCalls` assumes that polymerase slippage only occurs in STRs of 8 bases or more and only results in insertions or deletions of a single repeat unit. The likelihood of $a$ alt reads out of $d$ total reads in the case of a real somatic variant is given by the allele fraction clustering model. The likelihood in the case of polymerase slippage is the marginal of binomial likelihoods over a slippage rate with a uniform prior from 0 to 0.1, which is a regularized Beta function. Given priors and likelihoods, the error probability follows.

##### I. Normal Artifacts

A matched normal is useful not only for detecting germline variants but also for distinguishing technical artifacts for somatic mutations. Even a large panel of normals may not include, for example, mapping errors due to a rare SNP in a centromere, while a matched normal will exhibit such errors. `Mutect2` emits a normal artifact log odds annotation by applying the somatic likelihoods model to the normal. We combine this likelihood of the reads given that an artifact appears in the normal with a prior probability for normal artifacts to obtain a posterior error probability. Since this prior is unknown, we use the rate of technical artifacts in the tumor detected by `FilterMutectCalls` as a proxy.

##### J. Bad Haplotypes

`Mutect2` emits phasing information for calls in the same assembly region. We assign a "bad haplotype" probability equal to the greatest technical artifact probability of any in-phas call within a certain distance, by default 100 bases.

#### IV. LOCAL ASSEMBLY IN THE GATK

The GATK tools `HaplotypeCaller` and `Mutect2` assemble reads aligned within a window of several hundred base pairs into an assembly graph of local variation. Local haplotypes correspond to paths in this graph, and the downstream likelihood calculations in both tools involve aligning reads to these haplotypes. Here we describe how this graph is generated.

##### A. Correcting Reads

Read error correction is turned off by default in `HaplotypeCaller` and `Mutect2` and we are not familiar with it, nor have we validated it. Nonetheless, the code exists and can be turned on. The idea is as follows: large kmers are good because they contain more phasing information and therefore yield a simpler assembly graph. For example, two SNVs 20 bases apart with sequenced with error-free reads yield a graph with $2 \times 2 = 4$ paths if $k = 10$ but only 2 paths if $k = 40$, because the latter spans the phased SNVs. For example, consider a reference sequence TGAAACGTATTTGGG and an alt sequence with two phased SNVs, TGAAA(C→T)GTA(T→C)TTGGG. If $k = 3$ no kmer spans both SNVs and so the assembly graph containing two bubbles, one for each SNV, as in Figure 2. If $k = 5$, a kmer spans both SNVs and thus only two paths exist in the assembly graph, as in Figure 3.

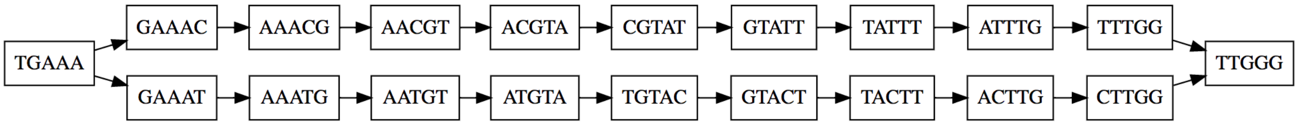


FIG. 3: Two haplotypes yielding assembly graph with a single bubble when $k$ is sufficient to phase variants.

One drawback is that larger kmers are more liable to be lost due to sequencing error simply because they contain more bases[7]. The idea of read error correction is to rescue kmers from the occasional error.

First, the `ReadErrorCorrector` kmerizes every read and counts the occurrences of each kmer. Then it builds a map from kmers to their corrected versions where kmers that appear often[8] map to themselves and kmers that appear only once[9] map to their nearest neighbor in Hamming distance[10] within a maximum of two mismatches[11]. Kmers that appear between 1 and 20 times are not part of the correction map.

Then, for every base in every read, the `ReadErrorCorrector` queries the kmer correction map for each overlapping kmer. For example, to correct the fourth base of read ACGTATTC if k = 3, it looks at the corrections for CGT, GTA, and TAT at their third, second, and first positions, respectively. If and only if the corrections are unanimous, the base is corrected. Note that corrected reads are used only for assembly and the original reads are used downstream.

#### B. Building the graph

Before we discuss how to construct the graph, let us confess a slight abuse of terminology. We refer to our assembly graph as a de Bruijn graph, that is, a graph in which the vertices are kmers and two kmers have a directed edge whenever they appear consecutively in a read. This is almost true (until the last part of assembly when we convert the graph into a more compact sequence graph), but there are a few differences:

- The edges of our graph record their multiplicity, the number of times the kmers they connect occur consecutively.
- The kmers of our graph come not just from reads but also the reference haplotype and, in GGA mode (see below), haplotypes generates from the given alleles.
- A non-unique kmer (see below) may be associated with multiple vertices.

Next, the `ReadThreadingAssembler` assembles the (corrected) reads over several different kmer sizes specified by the `kmerSize` argument[12]. If the given kmer sizes fail to produce a graph without cycles, or if more than 1/5 of the kmers in the graph are not unique in the sequences from which they come[13], the `ReadThreadingAssembler` repeatedly increases $k$ by 10 bases, starting from the largest initial $k$, until assembly succeeds or until $k$ has been increased 6 times, that is, by 60 bases[14].

The first step is to create a set of sequences to be kmerized and put into the de Bruijn graph. These sequences are: the reference haplotype, any requested alleles given by the `alleles` argument in `HaplotypeCaller`'s "genotype given alleles" (GGA) mode[15], and maximal subsequences of reads with base quality at least 10 by default[16] and at

---

[7] Another drawback is that longer kmers yield more dangling heads and tails in the graph (see below), especially near the ends of baited regions in exome sequencing.

[8] By default, 20 times or more. This threshold is set by the `minObservationsForKmerToBeSolid` command line argument.

[9] This is a hardcoded threshold.

[10] For simplicity we consider only substitution errors, eg the distance between ACGGT and AGGTG is 3

[11] This is a hardcoded threshold.

[12] By default, 10 and 25. This size is a compromise and no single value is the best choice for all regions. Large kmers are more likely to be unique and to yield a graph with no cycles, which is especially important in low-complexity regions, but they are more susceptible to decreased sensitivity due to errors and low coverage.

[13] This uniqueness condition is waived on the last attempt when $k$ has been increased by 60 bases.

[14] These constants, 10 and 6, are hard-coded, but the attempts to increase $k$ can be disabled completely with the `dontIncreaseKmerSizesForCycles` argument.

[15] This mode is disabled in `Mutect2` and incompatible with the reference confidence mode of `HaplotypeCaller`, which is the recommended best practice. We document it here for completeness. Before assembly, the GATK engine creates GGA haplotypes by substituting the reference bases with the alt bases from the GGA vcf in the reference haplotype. These GGA haplotypes are treated as non-reference sequences, that is, as long reads. The engine does not induce all possible haplotypes obtained from different phasings of GGA alleles, although depending on the kmer size many such haplotypes may appear in the assembly graph.

[16] This is controlled by the `minBaseQualityScore` argument.

least one kmer long. That is, a read with 100 bases and a base of quality 7 at position 70 yields sequences of length 69, 30 if $k = 10$ and a single sequence of length 69 if $k = 40$. We do not use mate information; that is, a read and its mate yield completely independent sequences, nor do we later post-process the graph using mate information. This is a potential direction for improvement.

Before building the graph, the `ReadThreadingAssembler` finds the set of all kmers that appear more than once in the same sequence i.e. more than once in the reference or in any read. These are called "non-unique" kmers and a kmer that is not non-unique is called "unique." Then each sequence is kmerized starting from the first unique kmer[17], with the exception that the reference sequence begins at its first kmer, unique or not. When a new kmer appear a new vertex is added to the graph. When a unique kmer that is already on the graph appears an edge is added, or the multiplicity of an extant edge is incremented, from the preceding kmer to the current one. When a non-unique kmer appears and there is already an edge joining the previous kmer to it this edge's multiplicity is incremented, but if an edge does not already exist a completely new vertex and edge are created[18]. The advantage of creating a new vertex for a non-unique kmer rather than re-using an old vertex is that it avoids spurious cycles. Up to this point, we have strayed from a canonical de Bruijn graph only as noted above. Note that this is a directed graph of kmers from aligned reads (i.e. aligned to the forward strand of the reference), not a bidirected graph of unaligned reads that could come from either strand, with the complications of associating a kmer with its reverse complement.

##### C. Cleaning the Graph

Before deciding on candidate haplotypes, the assembler simplifies the graph with the following heuristics to remove spurious paths and to merge variant paths that diverge from the reference.

- pruning: The assembler finds all maximal non-branching subgraphs ("chains") and removes those that 1) do not share an edge with the reference path and 2) contain no edges with sufficient multiplicity[19] While the default multiplicity threshold of 2 is quite permissive, it *does* cause `Mutect2` to lose sensitivity for deletions occurring in a single read[20].

  There is a command line flag `--adaptive-pruning` to turn on an adaptive pruning algorithm that adjusts itself to both the local depth of coverage and the observed sequencing error rate and removes chains based on a likelihood score. The score of a chain is the maximum of a left score and a right score, where the score on left (right) end of the chain is the active region determination log likelihood from the `Mutect2Engine`, treating the first (last) edge of the chain as a potential variant reads and all other outgoing (incoming) edges of the first (last) vertex in the chain as ref reads. The adaptive algorithm does this in two passes, where the first pass is used to determine likely errors from which to determine an empirical guess of the error rate.

  The adaptive pruning option is extremely useful for samples with high coverage, such as mitochondria and targeted panels, and for samples with variable coverage, such as exomes and RNA.

- dangling tails: The assembler only outputs haplotypes that start and end with a reference kmer, so it attempts to rescue paths in the graph that do not. To rescue a "dangling tails" – a path that ends in a non-reference kmer vertex – the assembler first traverses the graph backwards from this vertex to a reference vertex. If during traversal it encounters a vertex with more than one incoming edge it gives up[21] It also gives up if it encounters a vertex with more than one outgoing edge, that is, if the path branches again after diverging from the reference[22]. Then it generates the Smith-Waterman alignment of the branching path versus the reference path after the vertex at which they diverge. If the alignment's CIGAR contains three or fewer elements, that is, if the alignment has at most one indel, the assembly engine attempts to merge the dangling tail back into the reference.

  To merge the dangling tail back into the reference path, the assembler finds the beginning of the maximal common suffix of the dangling path and the reference path, that is, the point at which the sequences coverges[23]

---

[17] Note that a kmer that occurs only once in this particular sequence but more than once in some other sequence in non-unique.

[18] The fact that a non-unique kmer may be associated with multiple vertices on the graph is an important difference between this and a pure de Bruijn graph.

[19] By default 2. This is controlled by the `minPruning` argument.

[20] While a SNV occurring on a single read would not yield a confident somatic variant call, a long deletion in a non-STR context could easily be supported by a single read be due to the tiny probability of its arising from sequencing error.

[21] as opposed to doing eg depth-first search of all possible paths back to the reference.

[22] It seems like this could be changed to increase sensitivity.

[23] this is *not* where the *paths in the graph* converge (they don't) because kmers in the suffix disagree with the ref at upstream bases.

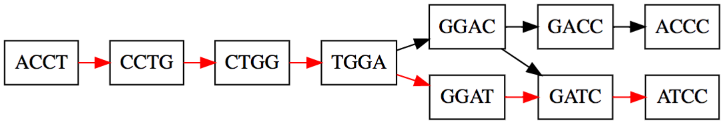


FIG. 4: A dangling tail merged back into the reference path. Reference path edges given by red arrows.

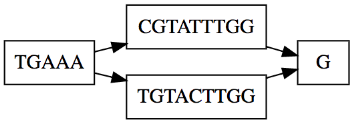


FIG. 5: A zipped sequence graph.

and adds an edge between the dangling path's vertex and the reference path's vertex at this position. This means that the graph is no longer a valid de Bruijn graph because the dangling vertex kmer and its succeeding reference vertex kmer do not overlap by $k-1$ bases. Nonetheless, this graph yields valid haplotypes when we later "zip" the graph's chains (see below) by accumulating the last base of each kmer.

Figure 4 shows the result of the tail from alt path ACCTGGA(T→C)CC being merged back into the reference path.

- dangling heads: This case and its treatment in the assembly engine is the mirror image of dangling tails.
- non-reference paths: After attempting to merge dangling heads and tails into the reference path, the assembler deletes edges and vertices belonging to dead ends i.e. non-branching subgraphs that start at non-reference source vertices or end at non-reference sink vertices. This is achieved by performing a breadth-first search of vertices moving forward from the reference source and a breadth-first search of vertices moving backwards from the reference sink and keeping only vertices found in both searches.
- zipping chains: A de Bruijn graph of kmers is convenient for performing assembly but is inefficient for summarizing the results of assembly. In this step the assembler converts the de Bruijn graph[24] into a sequence graph by combining all vertices in each maximal non-branching subgraph (i.e. each linear chain) into a single vertex containing the sequence implied by those vertices' kmers[25]. Thus, for example, a de Bruijn graph containing one reference path and one bubble for a single variant, such as the graph of Figure 3 yields a sequence graph with four vertices: the reference subgraphs before and after the bubble and the two sides of the bubble, as in Figure 5.
- merging diamonds: The assembler looks for nodes $A$ and $C$ such that multiple paths $A \to B_i \to C$ exist and absorbs the common prefix of $\{B_i\}$ into $A$ and the common suffix of $\{B_i\}$ into $C$. For example, if $k = 10$ and there was a single SNV bubble in the de Bruijn graph, the zipped sequence graph has a reference source path ($A$), a reference sink path ($C$) and two sides of the bubble ($B_1$ and $B_2$) that are each 10 bases long. Since $B_1$ and $B_2$ differ only in a single base, their common bases can be absorbed into $A$ and $C$ such that $B_1$ and $B_2$ contain only a single base each. Merging tails works the same way without the terminal vertex $C$ and merging

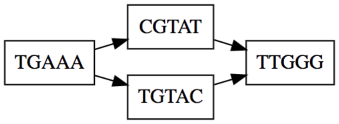


FIG. 6: A merged diamond.

[24] or rather, if dangling paths have been merged, an *almost* de Bruijn graph.
[25] That is, the concatenation of the last bases of all kmers, except for the first vertex which contributes its entire kmer if and only if it is a source.

common suffixes works the same way without the terminal vertex $A$. Figure 6 shows the graph of Figure 5 after the common suffix GGTT of the two sides of the bubble are absorbed into the terminal vertex.

- splitting common prefixes and suffixes: If there are multiple nodes $S_i$ with a common predecessor and successor of the form $S_i = A + x_i + B$, where $A$ is a common prefix, $B$ is a common suffix, and $x_i$ is unique to $S_i$, the assembler splits each $S_i$ such that $A$ and $B$ become independent nodes, with the $x_i$ in between.
- The clean-up steps of zipping chains, merging diamonds, tails, and common suffixes, and splitting common prefixes and suffixes are repeated until no more transformations occur.

When the `debugGraphTransformations` argument is used, the assembly engine emits graphviz .dot files for each of the steps above.

##### D. Finding Haplotypes

At this point there is one cleaned sequence graph for each successful kmer size. For each graph we find the best haplotypes[26] according the following score: the score of a path (haplotype) in a sequence graph is the sum over all branching vertices in the path of the log of the multiplicity of the outgoing edge in the path minus the log of the total multiplicity of all outgoing edges. This formula is implemented as a recursive depth-first search, sped up via dynamic programming: Starting from the reference source vertex, the haplotype finder adds branching scores and instantiates new haplotype finders for each edge whenever a path diverges and otherwise simply moves to the succeeding vertex without changing the accumulated score. The recursion ends when a haplotype finder reaches a sink vertex. The dynamic programming optimization is to cache the best sub-haplotypes found from every previously-reached vertex. Then, whenever this vertex is reached from a different prefix path, its best suffix paths are queried from the cache.

This description completely defines the best haplotypes; the actual implementation is a somewhat complicated recursive algorithm with lots of polymorphism.

#### V. PAIR HMM PROBABILISTIC REALIGNMENT IN HAPLOTYPECALLER AND MUTECT

After generating candidate haplotypes, the GATK tools `HaplotypeCaller` and `Mutect2` realign reads against these haplotypes to obtain a matrix of likelihoods for each read to be derived from each haplotype. Here we describe the probabilistic model specifying this likelihood as well as its computational implementation. We do not describe the translation of this implementation into native code optimized for vectorized architectures.

##### A. The Pair HMM model

We want to calculate the probability $P(\mathcal{R}|\mathcal{H})$ of read $\mathcal{R}$ to be sequenced from haplotype $\mathcal{H}$, where the haplotypes are sufficiently long that reads are contained within them. This likelihood is the sum of likelihoods of all possible alignments $\mathcal{A}$ of $\mathcal{R}$ to $\mathcal{H}$:

$$P(\mathcal{R}|\mathcal{H}) = \sum_{\mathcal{A}} P(\mathcal{R}, \mathcal{A}|\mathcal{H}) \tag{34}$$

We represent alignments as sequences

$$\text{alignment} = \{(i_1, j_1, s_1), (i_2, j_2, s_2) \ldots (i_N, j_N, s_N)\}, \tag{35}$$

where $i$ and $j$ represent positions within the read and haplotype, respectively, and $s_n \in \{M, I, D\}$ represents the states of match, insertion, and deletion of the read relative to the haplotype. For example, an alignment $\{(1,10,M),(2,11,M),(3,12,M),(4,13,M),(4,14,D),(4,15,D),(5,16,M),(6,17,M),(7,17,I),(8,18,M)\}$ means that positions 1 - 4 of the read match positions 10 - 13 of the haplotype, followed by a two-base deletion (advancing from 13 to 15 in the haplotype without advancing in the read), followed by a match of read positions 5 -

---

[26] This is 128 by default and set by the `maxNumHaplotypesInPopulation` argument.

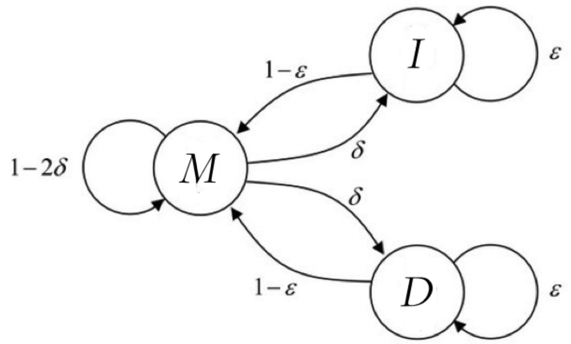


FIG. 7: Finite state machine of transitions among match, insertion, and deletion alignment states. In the absence of per-base BQSR indel qualities transition probabilities are parametrized by two constants, the indel start probability $\delta$ and the indel continuation probability $\epsilon$, such that $T_{MI} = T_{MD} = \delta$, $T_{MM} = 1-2\delta$, $T_{II} = T_{DD} = \epsilon$, $T_{IM} = T_{DM} = 1-\epsilon$, and $T_{ID} = T_{DI} = 0$.

6 with haplotype positions 16 - 17, followed by an insertion at read position 7, followed by a match. The allowable transitions $(i_n, j_n) \to (i_{n+1}, j_{n+1})$ are

- $(i, j, M/D/I) \to (i+1, j+1, M)$: match of read position $i+1$ with haplotype position $j+1$
- $(i, j, M/D) \to (i, j+1, D)$: deletion after read position $i$ – haplotype position $j+1$ is deleted
- $(i, j, M/I) \to (i+1, j, I)$: insertion at read position $i+1$

Note that the state label $s$ seems redundant because it can be reconstructed from the sequence of $i$ an $j$. While this is true, our model treats indel starts differently from indel continuations, and by including the state label we can distinguish these conveniently[27]. These transitions are illustrated as a finite state machine in Figure 7.

The read-alignment likelihood has two components. First is the probability of the sequence of match, insertion, and deletion states, which is

$$P(\mathcal{A}) = \prod_k T_{s_k, s_{k+1}}, \tag{36}$$

where $T$ is a matrix of state transition probabilities[28][29]. The index $k$ runs from 1 to the number of states in the alignment, that is, the read length plus the number of deleted reference bases. Next is the emission probability of the read bases given the haplotype bases they align to and the base qualities:

$$P(\mathcal{R}|\mathcal{A}, \mathcal{H}) = \prod_k P(r_{i_k}|h_{j_k}, q_{i_k})^{\mathrm{I}[s_k = M]}, \tag{37}$$

where $r_m$ and $h_n$ are the $m$th read bases and $n$th haplotype base and $q_m$ is the quality of the read's $m$th base. Note that this only includes alignments in the match state. The per-base emission is given directly from the definition of base quality:

$$P(b_2|b_1, q) = \begin{cases} \epsilon(q)/3 & (b_1 \neq b_2) \\ 1 - \epsilon(q) & (b_1 = b_2) \end{cases} \tag{38}$$

---

[27] That is, we can tell which type of transition it is by looking back one unit instead of two. Thus we have a first-order Markov model instead of a second-order Markov model.

[28] The gap continuation probability corresponds to a phred-scaled quality that is set by the `gcpHMM` argument, which is 10 by default, implying that $T_{DD} = T_{II} = 10^{-10/10} = 1/10$ and $T_{DM} = T_{IM} = 1 - T_{II}$. The indel start transitions are derived from the read's BQSR base insertion and base deletion qualities, if they exist, or a phred-scaled indel start quality of 45 if they do not. In production at the Broad Institute bams do not have BQSR indel qualities, hence the constant default is used. That is, $T_{MD} = T_{MI} = 10^{-4.5}$ and $T_{MM} = 1 - 2 \times 10^{-4.5}$.

[29] The elements of $T$ are either empirical or, (sometimes) in the case of indel start transitions, derived from BQSR. However in principle all elemnts of $T$ could be learned by applying the Baum-Welch algorithm to some known training data. For example, one could use a haploid cell line, for which any assembly region has a single haplotype.

where $\epsilon(q) = 10^{-q/10}$ is the error rate implied by the phred-scaled quality $q$.

#### B. Dynamic Programming

Define the matrices $M$, $I$ and $D$ by $M_{ij} =$ the total likelihood of *all* paths from the beginning of the read to position $i$ that end in a match state, and likewise for $D$ and $I$. Then the recursions

$$M_{ij} = P(r_i|h_j, q_i)\left(M_{i-1,j-1}T_{MM} + I_{i-1,j-1}T_{IM} + D_{i-1,j-1}T_{DM}\right) \tag{39}$$

$$I_{ij} = M_{i-1,j}T_{MI} + I_{i-1,j}T_{II} \tag{40}$$

$$D_{ij} = M_{i,j-1}T_{MD} + D_{i,j-1}T_{DD} \tag{41}$$

define the entire pair HMM algorithm:

**Algorithm 1** Pair HMM algorithm

Initialize $M_{0,j} = I_{0,j} = 0$ and $D_{0,j} = 2^{1020}/|\mathcal{H}|$ for $1 \leq j \leq |\mathcal{H}|$.
**for** $1 \leq i \leq |\mathcal{R}|$ **do**
    **for** $1 \leq j \leq |\mathcal{H}|$ **do**
        Calculate $M_{ij}$ via Equation 39.
        Calculate $I_{ij}$ via Equation 40.
        Calculate $D_{ij}$ via Equation 41.
    **end for**
**end for**
Total likelihood $P(\mathcal{R}|\mathcal{H})$ is $\sum_j (M_{\mathcal{R},j} + I_{\mathcal{R},j})$.

That the $i = 0$ rows of $M$ and $I$ are initialized to zero corresponds to starting at an imaginary position one base before the read start in a deletion state[30]. The factor of $1/|\mathcal{H}|$ corresponds to a flat prior on which $j$ an alignment starts at. This is important for a local alignment because we don't want to penalize reads that start in the middle of the haplotype. The curious factor of $2^{1020}$ is a huge number to prevent underflow – all multiplications are by numbers less than 1, so we needn't worry about overflow – which is a much more efficient approach than performing the computation in log space. The omission of $D$ from the returned value recognizes the fact that a terminal deletion is meaningless.

Finally, we note a shortcut that the GATK exploits: when two consecutive haplotypes of the same length[31] agree up to the $k$th position, the first $k$ columns of $M$, $D$ and $I$ are recycled and the inner loop is over $k < j \leq \mathcal{H}$.

[30] Since $T_{DI} = 0$ this means that an alignment may not start with an insertion, though it may begin with one. This limitation is unnecessary and could be fixed by initializing the $i = 0$ row of $I$ to a non-zero value as well.

[31] The condition on the same length could easily be relaxed simply by accounting for the constant $1/|\mathcal{H}|$ initialization.
